# Hawkish but helpful: When cultural group selection favors within-group aggression

**DOI:** 10.1101/001487

**Authors:** Ben Hanowell

## Abstract

The origin of cooperation is a central problem in evolutionary biology and social science. Cultural group selection and parochial altruism are popular but controversial evolutionary explanations for large-scale cooperation. Proponents of the cultural group selection hypothesis argue that the human tendency to conform—a consequence of our reliance on social learning—maintained sufficient between-group variation to allow group selection (which favors altruism) to overpower individual selection (which favors selfishness), whereupon large-scale altruism could emerge. Proponents of the parochial altruism hypothesis argue that altruism could emerge in tandem with hostility toward other groups if the combination of the two traits increased success in inter-group contests. Proponents of both hypotheses assume that cooperation is altruistic and that within-group conflict is antithetical to cooperation, implying that group selection for cooperation reduces within-group conflict. Yet within-group conflict need not be antithetical to cooperation. This essay uses a mathematical model to show that selection between groups can lead to greater within-group aggression if within-group aggression enhances the value of individually costly public goods contributions. This model may help to explain cross-cultural associations between warfare, socialization for aggression, aggressive sports, and interpersonal violence among humans. It may also apply to other forms of inter-group conflict among humans. Finally, the model suggests that group selection can lead to disharmony within groups, a caveat to the use of group selection models to inform social policy.

## 1.0 Introduction

Humans are a highly cooperative species (Smith 2010). Yet humans across cultures and throughout time have exhibited frequent within- and between-group aggression (Ember and Ember 1994; Ember and Ember 2007; Walker 2001). For these reasons, the origins, maintenance, and diversity of large-scale cooperation, within-group aggression, and inter-group conflict are central issues in evolutionary social science. In this essay, I draw on cross-cultural studies of warfare and aggression (Ember and Ember 1994; Ember and Ember 2007; Ross 1986; Russell 1972) and evolutionary game theory (Maynard Smith 1982) to motivate, derive, and analyze a simple model of coupled within-group cooperation and aggression. Contrary to two leading evolutionary hypotheses regarding large-scale cooperation and inter-group conflict, I show that selection between groups can in principle lead to greater within-group aggression than selection within groups if within-group aggression enhances the value of costly public goods contributions.

In the remainder of this section, I review two related hypotheses about the evolution of large-scale cooperation and inter-group conflict, and describe empirical evidence that refutes their key assumptions regarding the compatibility of within-group aggression with cooperation. This discussion motivates the formulation of a new evolutionary model. In section 2, I describe the classic evolutionary game theory models that combine to form the present model, derive the model’s payoff structure, analyze its stable states under selection within and between groups, and describe the conditions when selection between groups favors more interpersonal aggression than selection within groups. In section 3, I discuss the model’s results, limitations, applicability to human social behavior, and implications for recent attempts to use the group selection hypotheses to inform social policy.

### 1.1 The cultural group selection and parochial altruism hypotheses

Cultural group selection is a popular (Richerson and Boyd 2005; Wilson 2011) but controversial (Leigh Jr 2010; West et al. 2011) evolutionary explanation for large-scale cooperation among humans. Proponents of this hypothesis assume that large-scale cooperation entails individual altruism (in the biological sense; see West et al. 2007 for a definition), and that humans rely heavily on social learning that is biased toward conformity. Combining these assumptions, researchers argue that conformist social learning maintained sufficient between-group variation to allow group selection (which favors altruism) to overpower individual selection (which favors selfishness), whereupon large-scale altruism could emerge (Henrich 2004). Other researchers have proposed that altruism could have been maintained by genetic group selection alone (Bowles 2006).

Parochial altruism is another explanation for large-scale cooperation among humans. It is also an explanation for large-scale inter-group conflict. Citing experimental evidence (Bernhard et al. 2006) that humans are cooperative toward fellow group members but hostile (that is, parochial) toward outsiders, Choi and Bowles (2007) used a numerical simulation to argue that more altruistic and parochial groups could outperform more selfish and passive groups. García and van den Bergh (2011) confirmed and expanded these results with additional simulations. Proponents of the cultural group selection hypothesis have also formulated hypotheses for the emergence of ethnocentrism, which may underlie inter-group conflict (Boyd and Richerson 1985; Gil-White 2001). Unlike proponents of the cultural group selection hypothesis, proponents of the parochial altruism hypothesis argue that large-scale cooperation could have emerged solely through genetic group selection (Bowles 2009). Yet both hypotheses share the assumption that large-scale cooperation entails large-scale altruism. Thus they both emphasize selection between groups as the evolutionary mechanism that favors large-scale cooperation.

Both hypotheses also agree that prosocial norms (i.e., norms that encourage helping others for the sake of helping others) are the proximate mechanisms underlying large-scale cooperation (Bowles 2006; Henrich and Boyd 2001). Consequently, proponents of these hypotheses view between-group selection as a process that reduces antisocial behavior. One example of antisocial behavior is interpersonal aggression toward fellow group members. Wilson (1989: 267), referring to the classic hawk-dove game (Maynard Smith 1982) as a model for the evolution of aggression, noted that group selection leads to the disappearance of aggression, whereas individual selection allows for a stable mix of aggression and non-aggression. These arguments for the evolution of prosociality—combined with the apparent flexibility of human behavior and rapidity of cultural evolutionary processes (Boyd and Richerson 1985; Smith 2011), predict a negative cross-cultural association between warfare and interpersonal aggression within small-scale societies. This prediction, however, is at odds with the empirical evidence.

### 1.2 Frequent warfare is cross-culturally associated with greater within-group aggression

Ember and Ember (1994; 2007) used codes derived from the Human Relations Area Files to study the cross-cultural relationships between warfare, socialization for aggression in late boyhood, and frequency of interpersonal aggression within groups (specifically, assault and homicide) for small-scale societies. Combining path analysis with bivariate correlations, they argued convincingly that frequent warfare causes a society’s emphasis on socialization for aggression (not the other way around), and thus indirectly causes more interpersonal aggression within groups. Their analysis confirmed previous cross-cultural research that found links between within- and between-group conflict (Ross 1986; Russell 1972), and provided a window to understand the cross-cultural relationship between warfare and violent sports (Chick et al. 1997; Sipes 1973). More recently, Pinker (2011a; 2011b) has reviewed evidence that violence both within and between groups has declined over several millennia of human existence, which is circumstantial albeit longitudinal and comparative evidence for the coupling of within- and between-group conflict.

As for why warfare would causes socialization for aggression, Ember and Ember (1994; 2007) suggested that socialization for aggression prepares individuals for inter-group conflict, which poses both individual and group fitness benefits. One could argue further that experience with interpersonal aggression directly prepares individuals (thus groups) for combat. Yet interpersonal aggression (thus socialization for aggression) also poses individual and group costs by increasing morbidity and mortality while reducing group solidarity. Ember and Ember’s argument raises three important questions regarding cooperation and conflict. First, how would the evolutionary dynamics of large-scale cooperation in inter-group warfare interact with the evolutionary dynamics of interpersonal aggression within groups? Second, under what conditions do the evolutionary benefits of interpersonal aggression outweigh its costs? Finally, is it possible that selection between groups—perhaps facilitated by conformist social learning—would lead to more within-group aggression, an antisocial behavior, than individual selection? In the following sections, I derive and analyze an evolutionary game theoretic model to address these questions.

## 2.0 The hawkish cooperation model

### 2.1 Model building blocks: the hawk-dove and public goods games

The model in the following sub-sections combines two classic models from evolutionary game theory: the hawk-dove game and public goods games. The hawk-dove game models the evolution of aggression. The public goods game models the evolution of altruism. In the hawk-dove game, individuals pair randomly to contest a resource valued at *v* > 0. Doves yield to hawks (resulting in payoffs *v* to hawk and zero to dove) but share the resource with one another (yielding payoff *v*/2 to dove). Two hawks will fight for the resource. A hawk loses the fight half the time at cost *c* > *v*, yielding payoff (*v* – *c*)/2 to hawk. A stable proportion *v/c* of hawks exists under selection within groups (Maynard Smith 1982). Selection between groups favors doves because fighting injuries decrease group average payoff (Wilson 1989). In the public goods game, a cooperator contributes a benefit *b* > 0 to the public good at personal cost *x*, where 0 < *x* < *b*. A shirker consumes the public good without contributing. Under selection within groups, shirkers take over because cooperation is altruistic. Under selection between groups, cooperators take over because cooperation increases group average payoff. Both games have continuous strategy analogues, which produce mathematically equivalent results. See McElreath and Boyd (2007) for accessible reviews of the hawk-dove and public goods games. Because it combines these two games, I call the present model the “hawkish cooperation model.”

### 2.2 Payoff structure of the hawkish cooperation model

Imagine a large population of individuals partitioned into many large groups. Hawkish cooperators (HC) behave as hawks in within-group dyads, but as cooperators in a group-wide public goods game, which involves costly contribution to inter-group contests. Conversely, dovish shirkers (DS) behave as doves in within-group dyads, and do not contribute to the public good. Figure 1 depicts the structure of this model. The aggression of HC trains it for inter-group competition, increasing the public good by *a* ≥ 0 per HC. Yet HC aggression also decreases group solidarity, diminishing the public good by *s* ≥ 0 per HC. Fight injuries reduce HC efficiency in inter-group contests by *fc* per injury, which occurs with probability *h*^2^/2, and where *f* ≥ 0 scales the effect of injury on HC efficiency. Fighting and cooperation are costly (i.e., 0 < *v* < *c* and *x* > 0). I assume large groups due to our interest in large-scale social behavior; thus I ignore the fitness effect of an individual’s own public good contribution. Assuming very many groups, I ignore genetic drift. Thus the following payoffs *U* to HC and DS:

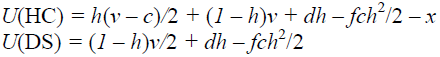

**Figure 1.** Schematic of the hawkish cooperation model. Individuals (dark ovals) play a hawk-dove game with a randomly selected partner (ovals containing two individuals) and a public goods game with all members of their group (ovals containing multiple dyads), which influences the outcome of between-group contests (bidirectional arrow connecting two groups).

Above, *h* is the proportion of HC and *d* = *a* - *s* is the benefit per HC to the public good, net the cost to group solidarity. If *d* ≤ 0, then the negative effect per HC on group solidarity cancels the positive effect on group preparedness. If *d* > 0, then the positive effect of one HC on group preparedness outweighs the negative effect on group solidarity. The overall effect of hawkish cooperation on the public good is ∂(*dh* − *fch*^2^/2)/*∂h* = *d − fch*; thus the public good is greatest where *h* = *d(fc)*^−1^. If *d* > *fc*, then any proportion of HC generates a public good. If *d* < 0, then hawkish cooperators are destructive. If *d* = 0, then HC are either destructive (if *f* > 0) or neutral (if *f* = 0).

### 2.3 Stable states of selection within groups

From selection within groups, two possible stable states emerge. If the payoff difference between a rare hawk and a common dove in the hawk-dove game exceeds the cost of altruism (i.e. if *v*/2 > *x*), then a stable internal equilibrium emerges where the proportion of HC is (*v* − 2*x*)/*c*. This equilibrium proportion of HC—which prevents fixation of HC (because *v < c* and *x* > 0)—increases with the value *v* of the contested resource and decreases with the costs of cooperation (*x*) and fighting (*c*). In this within-group selection scenario, unlike the classic public goods game, cooperators persist due to the individual benefit of sufficiently rare hawkishness (i.e., cooperation is not necessarily altruistic in this game). DS fixates instead if the cost of cooperation at least equals the payoff difference between a rare hawk and a common dove (i.e., if hawkish cooperation is altruistic). In this within-group selection scenario, unlike the classic hawk-dove game, hawks cannot invade doves. Appendix A derives these stable states.

### 2.4 Stable states of selection between groups

For selection between groups, a stable interior equilibrium exists where the proportion *h* of HC is (*d* − *x*)/(*c*(1 + *f*)), which is the proportion of HC that maximizes group average payoff. This proportion increases with the net benefit *d* per HC to the public good, but decreases with the cost *x* of altruism, the cost *c* of fighting, and the detrimental effect *f* of fighting injuries on HC efficiency. If the net benefit of an HC to the public good is greater than the average costs of cooperation in a group of all HC (i.e., if *d* > *x* + *c*(1 + *f*), then the equilibrium proportion *h* is unity. If the net benefit of an HC to the public good is less than the cost of cooperation (i.e., 0 < *d* ≤ *x*), then DS effectively fixates because the cost of cooperation cancels the net benefit of an HC to the public good. If HC is destructive and injuries decrease HC efficiency in inter-group contests (i.e., if *d* ≤ 0 < *x* and *f* > 0), then DS fixates because the overall effect of hawkish cooperation on the public good is negative. Unlike between-group selection in the classic hawk-dove game, doves might not fixate because the public good disappears without interpersonal aggression. Also unlike the classic public goods game, cooperators might not fixate because hawkishness is costly to groups with sufficiently common HC. Appendix B derives this stable state.

### 2.5 When selection between groups favors more aggression than selection within groups

Selection between groups may lead to greater interpersonal aggression than selection within groups only if the net benefit per HC to the public good is greater than the cost of cooperation (i.e., if *d* > *x*). Otherwise, the equilibrium proportion of HC arising from selection between groups is zero, which is less than or equal to that arising from selection within groups. If selection within groups favors DS, then selection between groups will lead to more interpersonal aggression than selection within groups if the net benefit per HC to the public good is greater than the cost of cooperation (i.e., *d* > *x*). Yet if selection within groups favors a stable polymorphism, then selection between groups will lead to greater interpersonal aggression only if the stable proportion arising from selection between groups (i.e., (*d* − *x*)/(*c*(1 + *f*))) exceeds that arising from selection between groups (i.e., (*v* − 2*x*)/*c*, which occurs if the following inequality holds:

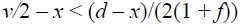

This inequality holds if the net benefit *d* per HC to the public good is sufficiently large relative to both the scale *f* of the effect of injuries on HC efficiency, and the payoff difference *v*/2 between a rare hawk and a common dove in the hawk-dove game. So although selection between groups may promote greater prosocial behavior in the form of greater contribution to the public good, it may also promote more interpersonal aggression, an antisocial behavior. Appendix C derives the conditions when selection between groups leads to more within-group aggression than selection within groups. In Appendix D, I describe a hawkish cooperation model where the strategy under selection is the probability that an individual plays HC, and show that the stable states of this model are mathematically equivalent to those in the discrete strategy model. Figure 2 illustrates the conditions when selection between groups favors more interpersonal aggression than selection within groups.

**Figure 2.** Average payoff *E*(*U*) as a function of the proportion *h* of hawkish cooperators (HC) in the group. Points A occur where the proportion of HC maximizes average payoff at *h* = (*d* − *x*)/*c*(1 + *f*) > 0. Point B occurs where individual selection (but not group selection) favors dovish shirkers (DS). Points C occur at a non-trivial stable polymorphism under individual selection, where *h* = (*v* – 2*x*)/*c*. Point D occurs where both group and individual selection favor DS. Thick curves represent *E*(*U*) given *h*. Dashed lines reference average payoff and proportion of HC at points of interest. **a.** A tragedy of hawkish cooperation when group selection favors a non-zero proportion of HC, but individual selection favors DS; *v* = 0.08, *c* = 0.25, *x* = 0.9, *f* = 0.1, *d* = 1. **b.** A tragedy of hawkish cooperation when group selection favors a proportion of HC greater than a stable polymorphism under individual selection; *v* = 0.4, *c* = 2, *x* = 0.1, *f* = 0.2, *d* = 1. **c.** No tragedy of hawkish cooperation when the stable polymorphism under individual selection has more HC than that under group selection; *v* = 1.5, *c* = 2, *x* = 0.1, *f* = 0.7, *d* = 1. **d.** No tragedy when both group and individual selection favor DS; *v* = 0.7, *c* = 1, *x* = 1.1, *f* = 0.75, *d* = 1.

## 3.0 Discussion

The hawkish cooperation model illustrates that cooperation and within-group aggression are not mutually exclusive. It suggests that within-group aggression may emerge via individual or group selection. It also shows that selection between groups may favor more antisocial behavior than selection within groups. Here, I discuss the model’s limitations, applicability to human social behavior, and implications for recent attempts by proponents of the group selection hypotheses to inform social policy.

### 3.1 Model limitations, plus prospects for extension and generalization

The hawkish cooperation model would benefit from extension and generalization. For simplicity, it assumes a pleiotropic effect on cooperation and interpersonal aggression, which are likely separate traits. Preliminary analysis of a multivariate hawkish cooperation model with a discrete strategy, not shown here, suggests that individual selection would yield the equilibria of the classic hawk-dove and public goods games. Multivariate group selection dynamics are more complex, possessing two equilibria sets and nonlinear parameter effects on equilibrium values. Future research will focus on multivariate models, allowing for non-zero covariance between traits.

The model also assumes very large groups in a very large population, but outcomes may differ for small groups in a finite population (Traulsen et al. 2005). Furthermore, the equilibrium fighting frequency diminishes in hawk-dove-type games where contestants recognize asymmetries in resource endowment or competitive ability (Gintis 2007; Kokko et al. 2006). Asymmetries diminish fighting frequency, thus decreasing the public good in the hawkish cooperation model. In addition, the model is related to scenarios such as the snowdrift game, where cooperative benefits or costs depend on the frequency of cooperators (Doebeli et al. 2004). Future research should explore the relationship between such models and those presented here.

The model also does not explicitly consider competition among groups, yet this is a key to the motivation for the hawkish cooperation model. Future research should model the relationships among between-group competition, interpersonal aggression, and the public good in more detail. Finally, I have not explicitly modeled socialization for aggression, a key component of Ember and Ember’s (1994; 2007) hypothesis about the association between warfare and interpersonal aggression. If the model did consider socialization for aggression, it would need to account for the mechanism of socialization. Because socialization among humans involves parents, but also “cultural” parents, the model would also have to account for the inclusive fitness benefits of socializing offspring for aggression, and the dynamics of social learning. These extensions and others would help to clarify the conditions when selection between groups can result in more within-group aggression than selection within groups.

### 3.2 Applicability of the model to human social behaviors

Despite the model’s simplicity, it provides insight into links between cooperation and interpersonal aggression among humans. Ember and Ember’s (1994; 2007) and others’ (Ross 1986; Russell 1972) findings of a cross-cultural association between warfare and interpersonal aggression is consistent with the hawkish cooperation model. Furthermore, martial training often involves violence toward fellow group members, creating a tradeoff between group solidarity and personal risk of injury (even death) on the one hand, and battle readiness on the other (Ember and Ember 2007). Examples include the hunting of Helot peasants by Spartan warrior initiates in the rite of *crypteia* (Jeanmaire 1913; Köchly 1835), stick fighting among young *Nguni* males (Morrell and Carton 2012), and dueling in many cultures.

The hawkish cooperation model may also apply outside the context of warfare. For example, aggressive competition within a modern business firm may increase firm preparedness for competition with other firms even if intra-firm aggression compromises firm productivity due to infighting. Consequently, firms may be favored that encourage more within-firm competitiveness than appears rational for individual workers. The hawkish cooperation model shows that these and similar behaviors may emerge under individual selection. Yet selection at higher levels may explain the existence of hawkish yet cooperative behavior beyond individual optima due to the group benefits of interpersonal aggression.

### 3.3 Implications for using group selection hypotheses to inform social policy

Traditionally, researchers have used the cultural group selection and parochial altruism hypotheses to explain the existence of prosociality toward fellow group members. By showing that antisocial norms can emerge from selection between groups, the hawkish cooperation model adds to the list of caveats (see Lehmann et al. 2008; and West et al. 2011) to recent attempts by group selection hypothesis proponents to inform social policy. Some proponents of the group selection hypothesis argue that promoting healthy inter-group competition will reduce intra-group conflict (Sober and Wilson 1998; Wilson 2011). Yet if group selection was important in human evolutionary history, we must consider its downsides along with its upsides regarding social harmony. The upside is that group selection may support greater contribution to the public good. One downside that emerges from the parochial altruism hypothesis is that group selection for altruism may be tied to conflict between groups. With the hawkish cooperation model, I highlight another potential downside to group selection: evoking human groupishness may promote conflict within as well as between groups.

## Acknowledgments

Thanks to Eric Alden Smith and Paul Lovell Hooper for comments.

## Appendix

### A. Stable states of selection within groups in the discrete strategy hawkish cooperation model

From the strategy payoffs in section 2.2, I derive the conditions for each of the two strategies to increase in frequency when rare within a group. I begin with the invasion of common DS by rare HC. The payoff to a rare HC in large group of DS is *v* – *x*. The payoff to common DS is *v*/2. Rare HC will invade common DS when the payoff of a rare HC is greater than that of a common DS:

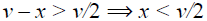

Conversely, common DS will resist invasion by rare HC when the payoff of common DS is at least that of rare HC:

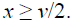

Similarly, rare DS will invade common HC when the payoff of rare DS exceeds that of common HC:

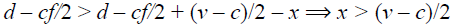

Note that this condition is always true because *v* < *c*, causing (*v* – *c*)/2 < 0 < *x*. For this reason, it is impossible for HC to resist invasion by DS. Together, these conditions allow two stable states, described in the main text.

### B. Stable states of selection between groups in the discrete strategy hawkish cooperation model

To find the proportion of HC that is stable under selection between groups, find the proportion of HC that maximizes average payoff by setting *∂E*(*U*)/∂*h* to 0 and solve for *h*. Then use the second derivative test ∂^2^*E*(*U*)/∂^2^*h* < 0 to double check if *E*(*U*) has a local maximum with respect to *h*. Since group average payoff is a concave parabolic function of the proportion of HC in the group, the function passes the second derivative test for a maximum. The proportion of HC that maximizes average payoff is presented in the main text. As discussed in the text, the maximum may exist outside the bounds of a valid proportion, leading to effective fixation of HC or DS.

### C. When selection between groups favors more aggression than selection within groups

The stable proportion of HC under selection between groups is (*d* − *x*)/(*c*(1 + *f*)), or zero if *d* < *x* (since a proportion cannot be less than zero). The stable proportion of HC under selection within groups is either zero (if *v*/2 < *x*) or (*v* − 2*x*)/*c* (if *v*/2 > *x*). Therefore, if *d* < *x*, selection between groups cannot lead to more interpersonal aggression than selection within groups because the equilibrium proportion of HC under selection between groups would be less than or equal to that under selection within groups. If, however, *d* > *x*, then selection between groups may lead to more interpersonal aggression than selection within groups. The necessary conditions for this to occur depend on which stable state obtains under selection within groups: fixation of DS, or a stable polymorphism.

If selection within groups leads to the fixation of DS, then selection between groups leads to more interpersonal aggression than selection within groups if the following inequality holds:

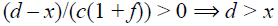

If selection within groups leads to a (non-trivial) stable polymorphism, then selection between groups leads to more interpersonal aggression than selection within groups if the following inequality holds:

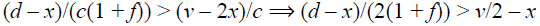

### D. Continuous strategy hawkish cooperation model

Suppose that a rare mutant with probability *p* = *q* + *δ* of playing HC emerges within a population where the expected probability of playing HC is *q*. Thus the following payoff to the mutant:

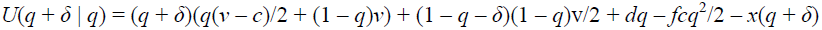

Here, *δ* describes the difference between the mutant and resident strategies. An equilibrium probability of playing exists under selection within groups where *∂U*(*q* + *δ* | *q*)/*∂q* = 0, which occurs where the resident strategy is *q* = (*v* – 2*x*)/*c*, the same as in the stable state under selection within groups in the discrete strategy model. To see if this equilibrium is stable, first determine the range of *q* for which selection favors lesser propensity to play HC:

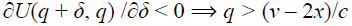

Selection favors lesser *q* if *q* is greater than the equilibrium. Next, determine the range of *q* for which selection favors greater propensity to play HC:

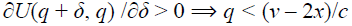

Selection favors greater *q* if *q* is lesser than the equilibrium. Therefore, a population perturbed a small amount from the equilibrium *q* in either direction will evolve back toward the equilibrium. Thus the equilibrium is stable and identical to that in the discrete strategy model.

To find the stable state under selection between groups, find the average propensity of playing HC that maximizes group average payoff. Suppose that a group has average propensity *y*. Thus the following group average payoff:

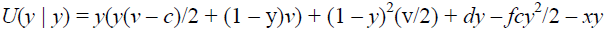

Take the derivative of the payoff function above with respect to y, set it to zero, and solve for y, which yields (*d* − *x*)/(*c*(1 + *f*)), identical to the stable state under selection between groups in the discrete strategy model. The second derivative is *−c*(1 + *f*), which is always negative, implying that (*d* − *x*)/(*c*(1 + *f*)) is a local maximum, and thus a stable state. Because the stable states under individual and group selection are equivalent to those in the discrete strategy model, the conditions for the tragedy of hawkish cooperation to exist are also equivalent.

